# Unified Genomic and Chemical Representations Enable Bidirectional Bio-synthetic Gene Cluster and Natural Product Retrieval

**DOI:** 10.1101/2025.05.31.656985

**Authors:** Guimei Liu, Yiting Li, Gabriel Ong, Fong Tian Wong, Dillon W. P. Tay, Yee Hwee Lim, Chuan Sheng Foo, Winston Koh

## Abstract

Natural product discovery is increasingly driven by the ability to analyze microbial genomes for biosynthetic gene clusters (BGCs) that encode secondary metabolites. While existing approaches have successfully linked BGCs to broad classes of chemical products, they typically operate in a single modality (genomic or chemical) limiting the scope of bidirectional prediction. In this work, we propose a multimodal framework that integrates genomic and chemical information by projecting embeddings derived from pretrained language models into a common representation space. We embed genomic sequences using a BGC foundation model and represent molecules through a chemical language model, then use a metric learning model to co-embed BGCs and their associated chemical structures. This co-embedding space allows us to quantify the similarity between BGCs and compounds using similarity measures, enabling both forward and inverse retrieval tasks. Beyond retrieval, we show that the shared space can guide strain selection for targeted compounds. By identifying BGCs closest to a query compound in the embedding space, we prioritize microbial strains that encode similar clusters, thereby streamlining genome mining and retrobiosynthetic design efforts. This approach represents a generalizable, scalable strategy to bridge biological and chemical modalities in natural product discovery.

## Introduction

The discovery of natural products increasingly relies on the sequencing of genomes to annotate and identify biosynthetic gene clusters (BGC) that produce bioactive compounds (***Scherlach and Hertweck, 2021***; ***Meesil et al., 2023***). Assigning a BGC sequence to its chemical product class is currently done by comparing sequence homology and domain architecture of a novel BGC to those of known BGCs with known products using Hidden Markov Model (HMM)-based tools (e.g. antiSMASH (***Medema et al., 2011***) and PRISM (***Skinnider et al., 2017***)), and more recently transformer-based models (e.g. DeepBGC (***Hannigan et al., 2019***) and BiGCARP (***Rios-Martinez et al., 2022***)). This has been very effective for annotation and broad classification of BGC sequences into major chemical classes of natural products (e.g. polyketides, saccharides, terpenes etc). However, there is an increasing need to address the inverse problem of working backwards from a target compound to select the appropriate BGC(s) capable of producing it or similar compounds, which is particularly important in the context of retrobiosynthesis. The ability to move in both directions (i.e. BGC to chemical, and chemical to BGC) could greatly increase the efficiency of searching and ranking microbial strains in large libraries where chemical product identity is often unknown. One way to achieve this would be multimodal projection of BGC sequences and chemical structures, which are two fundamentally disparate modes of information, into a common quantitative representation.

Recent advances in multimodal deep learning have revolutionized cross-modal search applications, spanning domains such as text-image retrieval (e.g., CLIP (***Radford et al., 2021***)) and biological function prediction through natural language embeddings (e.g., ProteinCLIP (***Wu et al., 2024***)). At the core of these breakthroughs is the use of vector embeddings derived from transformer-based models that encode diverse data types into unified numerical representations. Foundation models for BGCs, such as BiGCARP (***Rios-Martinez et al., 2022***) and DeepBGC (***Hannigan et al., 2019***), alongside chemical language models like ChemBERTa-2 (***Ahmad et al., 2022***), MoLFormer (***Ross et al., 2022***), and MolGPT (***Bagal et al., 2022***), illustrate the potential of these embeddings by capturing the structural and functional essence of their respective domains. These embeddings have shown utility in a variety of downstream tasks, from prediction of protein solubility (***Zhang et al., 2024***) in bioinformatics to prediction of bioactivity in cheminformatics (***Ross et al., 2022***).

Given the demonstrated richness of information captured by these foundation models, we reasoned that their embeddings, when projected in a common space, could reveal nuanced biochemical relationships between genomic and chemical modalities (***Keatinge-Clay, 2012; Oldfield and Lin, 2012***). To train such a model, we <u>B</u>GC<u>-C</u>hemical <u>Co-E</u>mbedding (BCCoE) that trains on paired examples of BGC sequences and their known smallmolecule outputs (Figure 1) from the curated MIBiG database (***Terlouw et al., 2023***).

**Figure 1.**
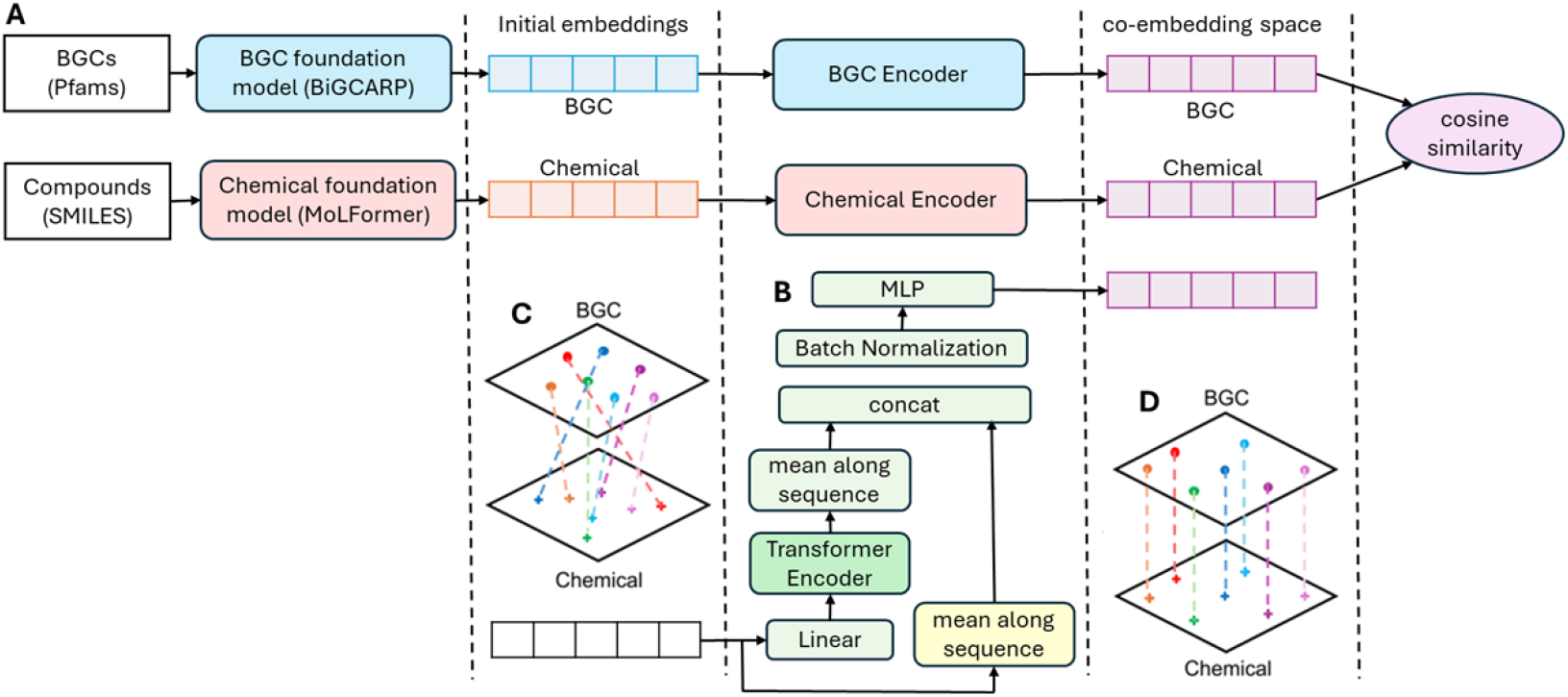
Architecture of our BCCoE model. **(A)** Overall model architecuture. **(B)** Encoder architecture. BGC encoder and chemical encoder have the same architecture, but they do not share model weights. **(C)** Initial embeddings of BGCs and compounds are not aligned. **(D)** Embeddings of BGCs and compounds from same pairs are well aligned and close to each other in the co-embedding space.

We hypothesize that the resulting common embedding space would produce a joint representation that effectively captures the genotype-chemotype relationship underlying BGC and their natural products. Such a shared space has the potential to facilitate seamless integration and retrieval across biological and chemical domains, enabling cross-modal analyses and downstream applications. One such application identified is bidirectional retrieval. Starting with a target compound, the joint embeddings could enable the identification, ranking, and prioritization of BGCs most likely to be responsible for its biosynthesis.

In a representative case study, we evaluated the BCCoE framework capability in inverting the natural product discovery process by starting from a known compound and identifying its likely biosynthetic strain of origin. We applied it to BE-54476-A/B, two anti-*Acinetobacter baumannii* tetramic acids originally isolated from Streptomyces sp. A58051 during a 54-strain screen (***Tay et al., 2024b***). Starting with the compounds’ SMILES, we generated chemical embeddings and projected them into the joint latent space. A cosine-similarity search against embedded MIBiG entries retrieved candidate BGCs. These candidate BGCs were then traced back to the strains in our 54-strain library via BLASTN search using the corresponding nucleotide sequences. The true producer strain was identified in the top 10 ranked strains in this workflow illustrating the utility of joint embeddings in sharpening strain prioritization.

## Results

### BGC-chemical co-embedding (BCCoE) for cross-modal alignment

We hypothesized that projecting biosynthetic gene clusters (BGCs) and their corresponding natural products into a shared latent space could reveal underlying biochemical relationships through spatial proximity. To evaluate this hypothesis, we developed a cross-modal representation learning framework, termed BGC-Chemical Co-Embedding (BCCoE), which aligns genomic and molecular information within a unified vector space (Figure 1). Our approach builds on recent advances in largescale pre-trained language models, which are capable of capturing complex structural and functional representations. Specifically, we employed BiGCARP (***Rios-Martinez et al., 2022***) to encode Pfam domain sequences of BGCs and MoLFormer (***Ross et al., 2022***) to generate embeddings of chemical SMILES strings. These transformer-based models provide informative, modality-specific embeddings; however, the resulting representations initially reside in distinct and unaligned spaces, as illustrated in Figure 1C. This necessitates an additional mapping strategy to bring the two modalities into correspondence.

To learn this alignment, we introduced modality-specific encoders—one for BGCs and one for compounds—each parameterized by a transformer architecture, and trained them using deep metric learning. We adopted the N-pair loss function (***Sohn, 2016***) as our optimization objective, which encourages known BGC-compound pairs to converge in the co-embedding space while simultaneously repelling non-associated pairs. Unlike binary cross-entropy or triplet loss, N-pair loss compares each positive example to multiple negatives sampled within the same mini-batch, resulting in stronger contrastive supervision and more robust alignment. Details of the model architecture and training procedure are described in methods section.

The resulting co-embedding space supports efficient bidirectional retrieval between BGCs and compounds, using cosine similarity as the similarity metric (Equation 1). In this approach, higher similarity scores indicate greater likelihood that a compound is the biosynthetic product of a given BGC, or conversely, that a BGC is capable of producing a queried compound. This enables systematic exploration of genomic–chemical relationships using simple nearest-neighbor queries (Figure 2). To assess the utility of this representation for natural product discovery, we formulated a retrieval task that reflects practical strain selection workflows. In the forward direction, the task involves ranking candidate compounds for a given query BGC based on embedding similarity. In the reverse direction, it involves ranking candidate BGCs for a given compound. Model performance was evaluated using four standard metrics computed at various cutoff thresholds: the number of true BGC-compound associations retrieved (hits at top-K), the fraction of known associations recovered (recall at top-K), the proportion of retrieved candidates that are correct (precision at top-K), and the improvement in retrieval relative to random ranking (lift at top-K). Together, these metrics quantify the extent to which the learned embedding space enriches for true biosynthetic relationships, supporting its application to hypothesis generation and prioritization in strain selection workflows.

**Figure 2.**
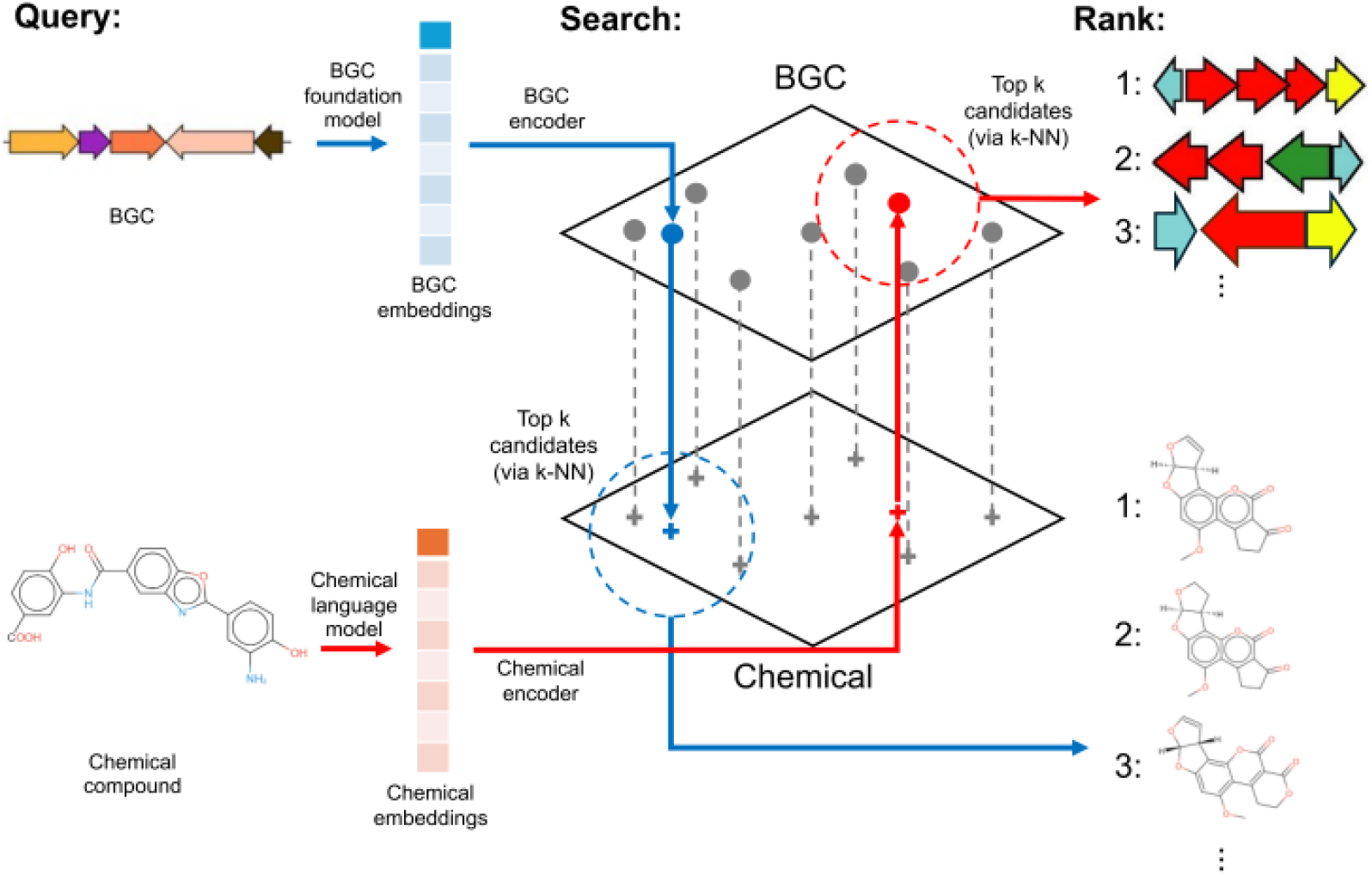
(A) Queries in the form of a novel BGC (top) or chemical compound (bottom) were embedded using their respective pre-trained foundation models. These embeddings were then projected by our trained BGC and chemical encoders into a common co-embedding space, where the nearest embeddings of the opposite modality (BGCs for chemical queries and chemicals for BGC queries) were retrieved using k-Nearest Neighbor search (k-NN), and ranked based on proximity.

### Cross-modal alignment enables accurate bidirectional retrieval

We evaluated the performance of our co-embedding framework across three settings designed to test its generalizability: (i) ten-fold cross-validation on all known BGC-compound pairs, (ii) class holdout evaluation, where all BGCs producing a specific compound class were excluded from training, and (iii) temporal generalization, where models trained on MIBiG version 3.1 were used to predict novel links in the subsequently released MIBiG version 4.0. These evaluation settings reflect different real-world deployment scenarios, including annotation of known compound classes, extrapolation to new classes, and forward prediction of unseen genomic-chemical links.

We compare our model with the no-alignment approach, which performs nearest neighbor search in the original embedding spaces directly without alignment. Two versions of the no-alignment approach are implemented: KNN and KNN-2hop, and they are described in methods section. Both of them use cosine similarity over the original embeddings generated by BGC and chemical foundation models. The difference between the two is that KNN uses either similarities between BGCs or similarities between compounds depending on the query while KNN-2hop uses both similarities in order to retrieve novel BGCs or compounds not in the training data.

In the 10-fold cross-validation setting, we randomly split the dataset into 10 folds either based on BGCs or compounds, and then use one fold as testing data and the remaining nine folds as training data. It represents a relatively ideal situation for machine learning where the training data and the testing data have similar distribution. The performance of the three models under this setting is shown in Figure 3 and exp 1 & 2 in Table 1 where the performance of the models is aggregated over the 10 folds. Our model with N-pair loss shows a very high enrichment compared with random guessing especially when K is small. In particular, when the task is to retrieve BGCs for given compounds, our model with N-pair loss can pick up around 58.9% of the ground-truths even at just top-5. In comparison, random guessing can only pick up 0.2% of the ground-truths at top-5.

**Table 1.** #hits, Recall and Precision of different models for bidirectional retrieval.

| exp | setting | top-K | random |  | KNN |  |  | KNN-2hop |  |  | BCCoE |  |  |
| --- | --- | --- | --- | --- | --- | --- | --- | --- | --- | --- | --- | --- | --- |
|  |  |  | #hits | Recall | #hits | Recall | Prec | #hits | Recall | Prec | #hits | Recall | Prec |
| 1 | 10-fold CV<br>BGC2Chem | 10 | 11.4 | 0.3% | 443 | 12.9% | 2.23% | 748 | 21.9% | 3.76% | 1125 | 32.9% | 5.66% |
|  |  | 100 | 114.7 | 3.4% | 521 | 15.2% | 0.26% | 1422 | 41.6% | 0.72% | 2243 | 65.5% | 1.13% |
| 2 | 10-fold CV<br>Chem2BGC | 10 | 12.6 | 0.4% | 1836 | 53.7% | 6.14% | 2075 | 60.6% | 6.94% | 2235 | 65.3% | 7.47% |
|  |  | 100 | 128.5 | 3.8% | 1900 | 55.5% | 0.64% | 2431 | 71.0% | 0.81% | 2864 | 83.7% | 0.96% |
| 3 | BGC classes<br>BGC2Chem | 10 | 14.9 | 0.3% | 60 | 1.4% | 0.25% | 145 | 3.4% | 0.59% | 253 | 5.9% | 1.04% |
|  |  | 100 | 145.6 | 3.4% | 72 | 1.7% | 0.03% | 478 | 11.1% | 0.20% | 1063 | 24.6% | 0.44% |
| 4 | BGC classes<br>Chem2BGC | 10 | 16.5 | 0.4% | 25 | 0.6% | 0.07% | 176 | 4.0% | 0.46% | 333 | 7.5% | 0.87% |
|  |  | 100 | 166.1 | 3.8% | 28 | 0.6% | 0.01% | 502 | 11.3% | 0.13% | 1333 | 30.1% | 0.35% |
| 5 | 3.1→4.0<br>BGC2Chem3.1 | 10 | 0.1 | 0.0% | 14 | 3.0% | 0.71% | 16 | 3.4% | 0.82% | 23 | 4.9% | 1.17% |
|  |  | 100 | 1.0 | 0.2% | 21 | 4.4% | 0.11% | 24 | 5.1% | 0.12% | 25 | 5.3% | 0.13% |
| 6 | 3.1→4.0<br>Chem2BGC3.1 | 10 | 1.9 | 0.4% | 127 | 26.8% | 2.78% | 168 | 35.5% | 3.68% | 188 | 39.7% | 4.11% |
|  |  | 100 | 17.6 | 3.7% | 146 | 30.9% | 0.32% | 223 | 47.1% | 0.49% | 313 | 66.2% | 0.68% |
| 7 | 3.1→4.0<br>BGC2Chem4.0 | 10 | 1.7 | 0.4% | 14 | 3.0% | 0.71% | 126 | 26.6% | 6.43% | 180 | 38.1% | 9.18% |
|  |  | 100 | 15.7 | 3.3% | 21 | 4.4% | 0.11% | 230 | 48.6% | 1.17% | 329 | 69.6% | 1.68% |
| 8 | 3.1→4.0<br>Chem2BGC4.0 | 10 | 1.8 | 0.4% | 127 | 26.8% | 2.78% | 168 | 35.5% | 3.68% | 190 | 40.2% | 4.16% |
|  |  | 100 | 17.8 | 3.8% | 146 | 30.9% | 0.32% | 236 | 49.9% | 0.52% | 335 | 70.8% | 0.73% |

**Figure 3.**
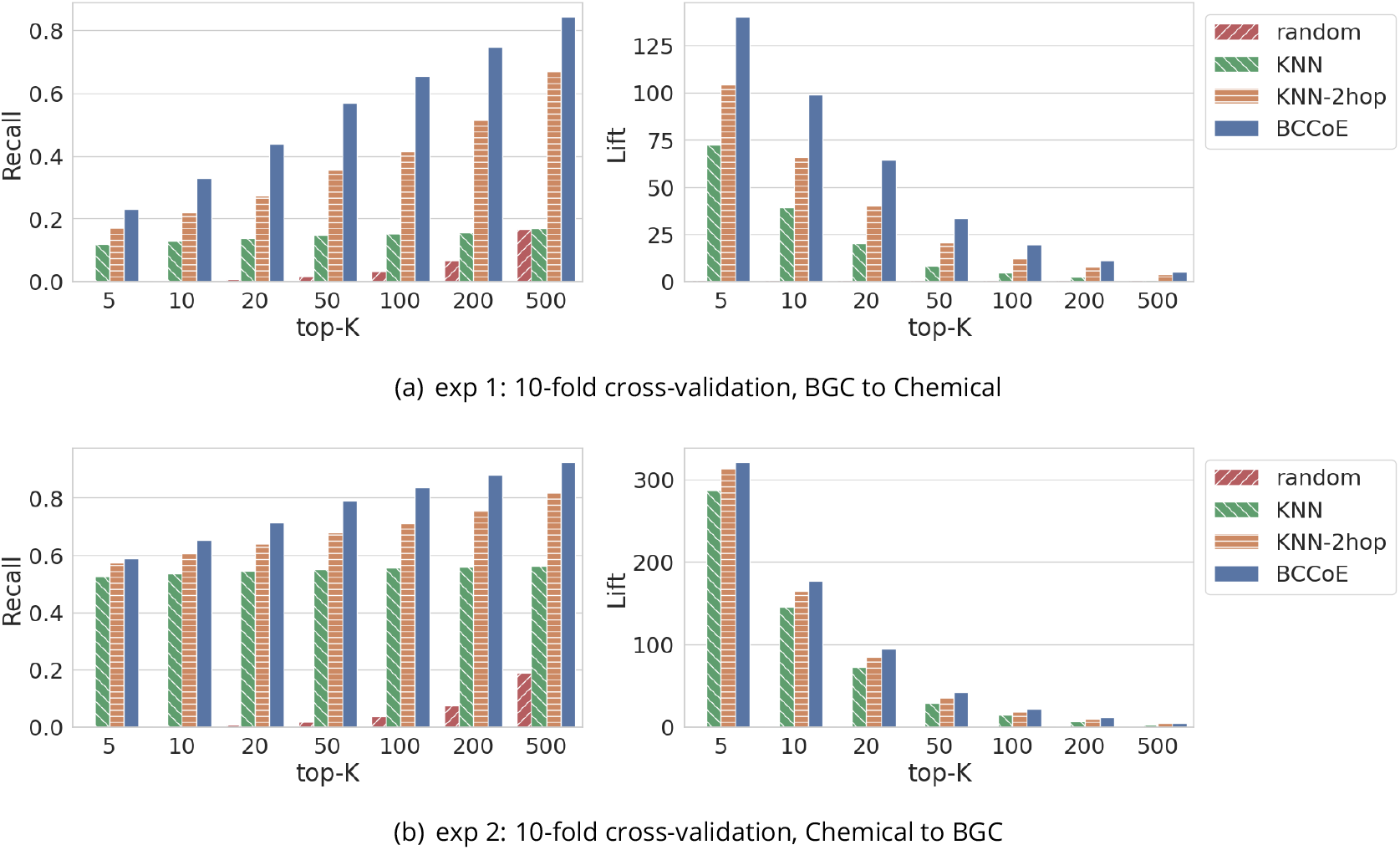
Model performance for bidirectional retrieval when using 10-fold cross-validation: Recall (left) and Lift (right)

Holding out one BGC product class splits data based on BGC product classes. BGCs or compounds in one specific BGC class are used as testing data and the remaining data are used as training data. This setup tests the ability for our model to transfer to novel BGC product classes; it represents a very challenging situation for machine learning because the training data and the testing data have very distinct distributions. The performance of our model under this setting is shown in Figure 4 and exp 3 & 4 in Table 1 where the performance of our model is aggregated over all the BGC product classes. Compared with using 10-fold cross-validation, the performance of our model drops significantly in these two experiments as expected. Nevertheless, our model using N-pair loss can still achieve a lift of 17.0 and 20.2 at top-10 over random guessing for the two retrieval tasks respectively, which is 74.5% and 89.2% better over KNN-2hop respectively. This results highlight the versatility of the representations in the co-embedding space in identifying meaningful associations between BGCs and chemicals, even in under-explored or newly discovered biological and chemical spaces.

**Figure 4.**
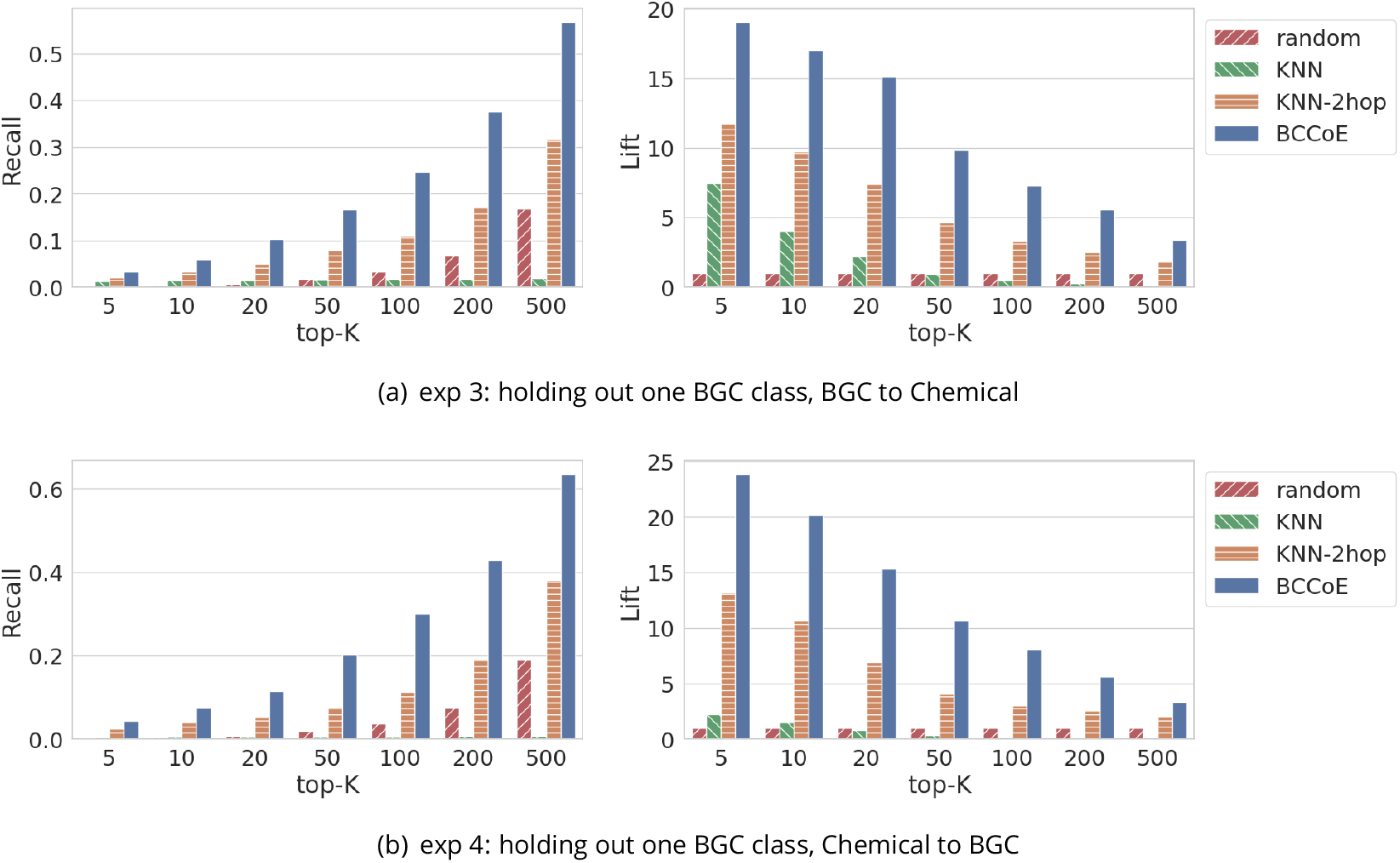
Model performance for bidirectional retrieval when holding out one product class: Recall (left) and Lift (right)

The performance of our model for predicting new pairs in MIBiG 4.0 using existing pairs in MIBiG 3.1 is shown in Figure 5 and exp 5-8 in Table 1. This setting represents a more realistic setting where we use known positive pairs to predict novel new pairs. There are 473 new BGC-compound pairs in MIBiG 4.0 compared with MIBiG 3.1 as shown in Table 3. Only 29 of them involving existing compounds in MIBiG 3.1 as shown in Table 5. Our BCCoE model is able to pick up 23 of them at top-10 when retrieving from existing compounds in MIBiG 3.1 (exp 5) while KNN-2hop retrieves only 16 of them at top-10. The number of hits of BCCoE at top-10 increases to 180 (38.1%) when retrieving from compounds from MIBiG 4.0 (exp 7). Most of the new pairs involve existing BGCs. BCCoE is able to pick up 188 (39.7%) new pairs at top-10 when retrieving from existing BGCs from MIBiG 3.1 (exp 6).

**Figure 5.**
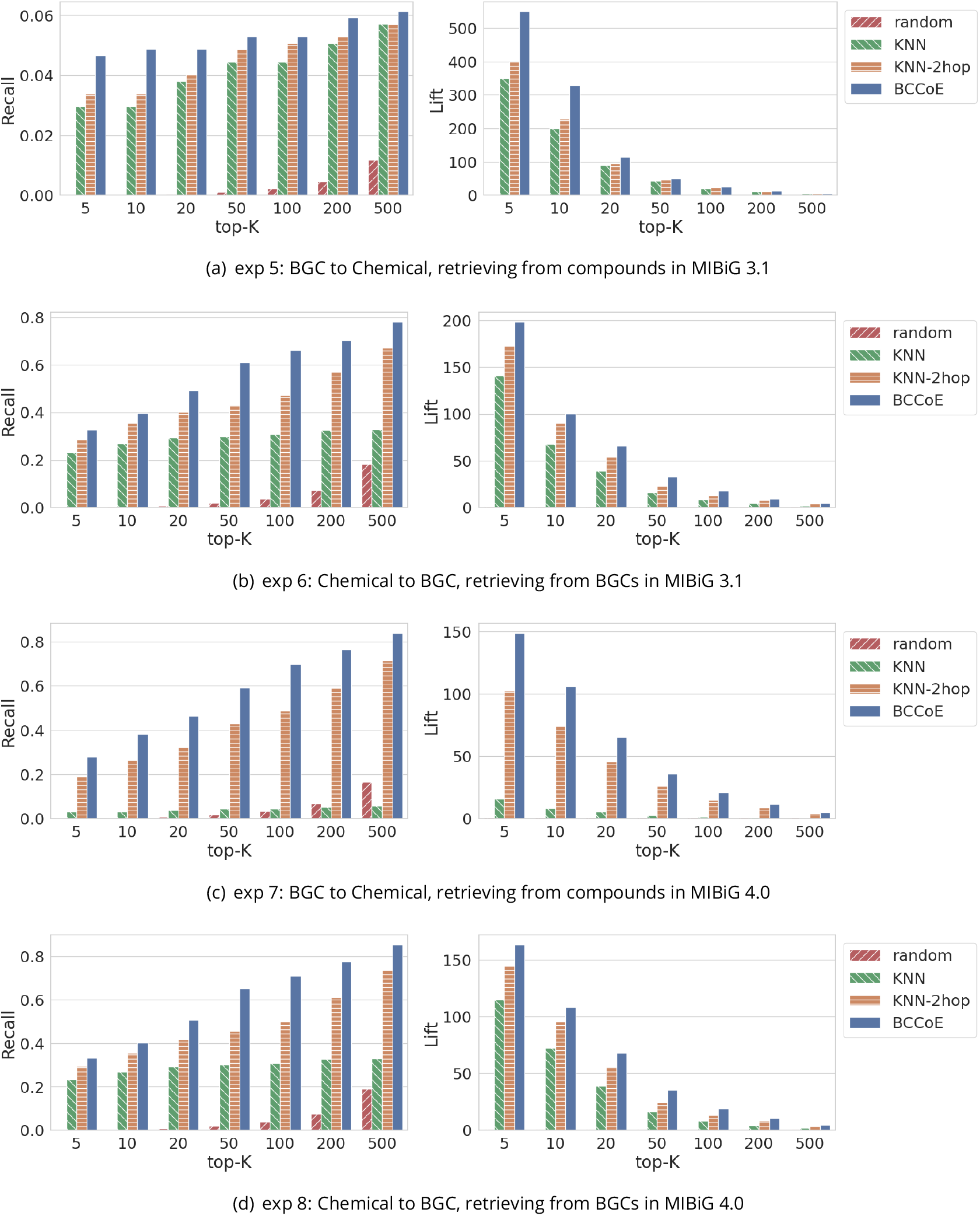
Model performance when using MIBiG 3.1 to predict new pairs in MIBiG 4.0: Recall (left) and Lift (right)

Together, these results demonstrate the robustness and utility of our co-embedding model in both well-studied and emerging discovery settings. Across all three evaluation paradigms, BCCoE consistently outperformed the unaligned embedding approaches, particularly at low K values, where precision and enrichment are critical for practical screening. These improvements suggest that aligning genomic and chemical representations in a shared latent space can substantially streamline strain prioritization, enabling more focused experimental validation and accelerating the discovery of novel natural products.

### Visualizing alignment in the learned co-embedding space

To qualitatively evaluate the extent of cross-modal alignment achieved by our co-embedding framework, we visualized the latent representations of BGCs and compounds using two-dimensional projections generated by t-distributed stochastic neighbor embedding (t-SNE) (***van der Maaten and Hinton, 2008***). The results are shown in Figure 6.

**Figure 6.**
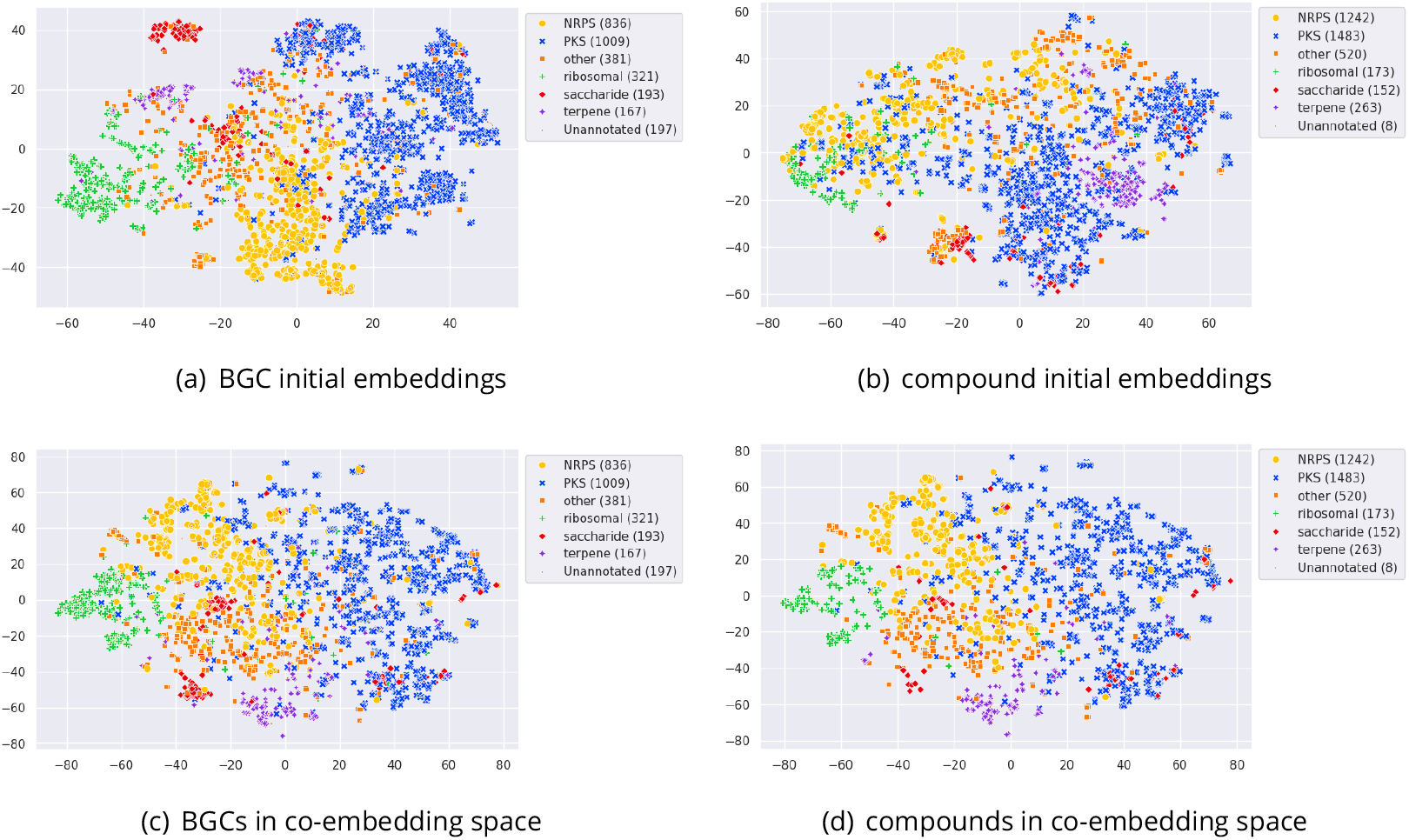
Visualization of BGCs and compounds: initial embeddings (top) and their embeddings in the co-embedding space (bottom). Numbers in parentheses are the number of BGCs/compounds in the product classes.

Prior to alignment, the initial embeddings produced by BiGCARP and MoLFormer displayed distinct clustering patterns within each modality. Both BGC and compound embeddings exhibited some grouping according to BGC product classes, indicating that the foundational models capture meaningful biochemical signals from Pfam domain sequences and SMILES strings, respectively. However, the embeddings of BGCs and compounds occupied separate regions of the projection space, reflecting poor alignment between the two modalities and limiting their utility for direct comparison or retrieval.

Following training with our deep metric learning objective, the joint co-embedding space exhibited markedly improved cross-modal alignment. As shown in Figure 6 panels (c) and (d), BGCs and their corresponding compounds now occupy overlapping regions of the space, consistent with the model’s objective to co-locate known BGC-compound pairs. These aligned representations not only facilitate bidirectional retrieval—as quantitatively demonstrated in prior sections—but also exhibit structure consistent with chemical class, suggesting potential utility for clustering, classification, and exploratory analyses. This visualization reinforces the interpretability and coherence of the learned latent space, supporting its application to both supervised and unsupervised tasks in natural product discovery.

### Case study: Inverting Natural Product Discovery for BE-54476 Tetramic Acids

To evaluate the practical utility of our multimodal co-embedding framework in real-world natural product discovery, we applied it retrospectively to the case of BE-54476-A and -B, two tetramic acid compounds with anti-*Acinetobacter baumannii* activity. These molecules were originally isolated from Streptomyces sp. A58051 during a multi-strain screening campaign involving 54 actinomycete genomes (***Tay et al., 2024b***).

We began by embedding the SMILES representations of BE-54476-A and -B using our pre-trained chemical encoder and projecting them into the learned joint chemical–genomic co-embedding space. From there, we performed cosine similarity search over the BGCs in MIBiG 4.0 to retrieve the top-ranked candidate biosynthetic gene clusters. Each retrieved MIBiG BGC was then mapped to our in-house strain collection by extracting its amino acid sequence and running tblastn against all 54 assembled genomes. For each strain, we computed a ranking score based on the number of significant alignments to the candidate BGCs (Equation 13).

To benchmark the effectiveness of the co-embedding approach, we compared it against KNN-2hop. KNN-2hop first finds similar chemicals in the unaligned chemical embedding space and then transitions to the unaligned genomic embeddings via the BGC-compound pairs in MIBiG 4.0. Both methods successfully identified Streptomyces sp. A58051—the true producing strain—within the top 5 ranked BGCs. When considering only the top 1 BGC for both BE-54476-A and -B, the coembedding approach ranked A58051 at position 4 out of the 54 strains, with 93 high-scoring tblastn hits, while KNN-2hop ranked it at position 6 with 77 hits (Table 2). These findings demonstrate that cross-modal alignment sharpens retrieval precision and streamlines strain prioritization, offering a path toward inverting the natural product discovery process.

**Table 2.** Top 5 candidate BGCs for BE-54476-A/B determined by BCCoE and KNN-2hop, and their corresponding strain hits. Correct strain hits are highlighted in yellow.

| Compound | Method | BGC | Score | Top 1<br>Strain | Top 2<br>Strain | Top 3<br>Strain | Top 4<br>Strain | Top 5<br>Strain | Top 6<br>Strain | Top 7<br>Strain | Top 8<br>Strain | Top 9<br>Strain | Top 10<br>Strain |
| --- | --- | --- | --- | --- | --- | --- | --- | --- | --- | --- | --- | --- | --- |
| BE-54476-A | BCCoE | BGC0001738 | 0.67975897 | A5252<br>(149) | A80510<br>(136) | A5858<br>(109) | A58051<br>(93) | A11345<br>(93) | A61715<br>(88) | A8567<br>(70) | A2056<br>(68) | T1312<br>(64) | A6562<br>(61) |
|  |  | BGC0001255 | 0.6309823 | A5252<br>(160) | A80510<br>(147) | A5858<br>(110) | A61715<br>(97) | A11345<br>(82) | A58051<br>(77) | A8567<br>(77) | A2056<br>(66) | T1312<br>(63) | T354<br>(62) |
|  |  | BGC0003145 | 0.55855453 | A5252<br>(194) | A80510<br>(157) | A5858<br>(127) | A61715<br>(114) | A58051<br>(99) | A11345<br>(99) | A8567<br>(92) | A2056<br>(90) | A33995<br>(84) | A34001<br>(75) |
|  |  | BGC0002304 | 0.515368 | A5252<br>(158) | A80510<br>(153) | A5858<br>(123) | A11345<br>(102) | A61715<br>(101) | A58051<br>(99) | A2056<br>(86) | A6562<br>(79) | A8567<br>(76) | T354<br>(73) |
|  |  | BGC0002538 | 0.47208548 | A5252<br>(126) | A80510<br>(92) | A5858<br>(92) | A61715<br>(74) | T1312<br>(66) | A11345<br>(55) | A33995<br>(49) | A58051<br>(47) | A34001<br>(43) | A34053<br>(43) |
|  | KNN-2hop | BGC0001255 | 0.944217086 | A5252<br>(160) | A80510<br>(147) | A5858<br>(110) | A61715<br>(97) | A11345<br>(82) | A58051<br>(77) | A8567<br>(77) | A2056<br>(66) | T1312<br>(63) | T354<br>(62) |
|  |  | BGC0001738 | 0.94100678 | A5252<br>(149) | A80510<br>(136) | A5858<br>(109) | A58051<br>(93) | A11345<br>(93) | A61715<br>(88) | A8567<br>(70) | A2056<br>(68) | T1312<br>(64) | A6562<br>(61) |
|  |  | BGC0000685 | 0.931540132 | A8567<br>(46) | A5252<br>(45) | A80510<br>(43) | A2056<br>(40) | A5858<br>(39) | A11345<br>(35) | A30639<br>(31) | A1090<br>(30) | T108<br>(30) | T12<br>(30) |
|  |  | BGC0001449 | 0.920163807 | A5252<br>(206) | A80510<br>(180) | A5858<br>(156) | A61715<br>(130) | A58051<br>(125) | A11345<br>(123) | A2056<br>(116) | A8567<br>(108) | A6562<br>(102) | A34001<br>(101) |
|  |  | BGC0001881 | 0.919855788 | A5252<br>(225) | A80510<br>(177) | A5858<br>(162) | A61715<br>(135) | A11345<br>(132) | A58051<br>(125) | A2056<br>(115) | A8567<br>(111) | A6562<br>(103) | T1312<br>(97) |
| BE-54476-B | BCCoE | BGC0001738 | 0.7113037 | A5252<br>(149) | A80510<br>(136) | A5858<br>(109) | A58051<br>(93) | A11345<br>(93) | A61715<br>(88) | A8567<br>(70) | A2056<br>(68) | T1312<br>(64) | A6562<br>(61) |
|  |  | BGC0001255 | 0.6409487 | A5252<br>(160) | A80510<br>(147) | A5858<br>(110) | A61715<br>(97) | A11345<br>(82) | A58051<br>(77) | A8567<br>(77) | A2056<br>(66) | T1312<br>(63) | T354<br>(62) |
|  |  | BGC0003145 | 0.589077 | A5252<br>(194) | A80510<br>(157) | A5858<br>(127) | A61715<br>(114) | A58051<br>(99) | A11345<br>(99) | A8567<br>(92) | A2056<br>(90) | A33995<br>(84) | A34001<br>(75) |
|  |  | BGC0002304 | 0.54288 | A5252<br>(158) | A80510<br>(153) | A5858<br>(123) | A11345<br>(102) | A61715<br>(101) | A58051<br>(99) | A2056<br>(86) | A6562<br>(79) | A8567<br>(76) | T354<br>(73) |
|  |  | BGC0002538 | 0.5189768 | A5252<br>(126) | A80510<br>(92) | A5858<br>(92) | A61715<br>(74) | T1312<br>(66) | A11345<br>(55) | A33995<br>(49) | A58051<br>(47) | A34001<br>(43) | A34053<br>(43) |
|  | KNN-2hop | BGC0001255 | 0.946465492 | A5252<br>(160) | A80510<br>(147) | A5858<br>(110) | A61715<br>(97) | A11345<br>(82) | A58051<br>(77) | A8567<br>(77) | A2056<br>(66) | T1312<br>(63) | T354<br>(62) |
|  |  | BGC0001738 | 0.942865729 | A5252<br>(149) | A80510<br>(136) | A5858<br>(109) | A58051<br>(93) | A11345<br>(93) | A61715<br>(88) | A8567<br>(70) | A2056<br>(68) | T1312<br>(64) | A6562<br>(61) |
|  |  | BGC0000685 | 0.929388702 | A8567<br>(46) | A5252<br>(45) | A80510<br>(43) | A2056<br>(40) | A5858<br>(39) | A11345<br>(35) | A30639<br>(31) | A1090<br>(30) | T108<br>(30) | T12<br>(30) |
|  |  | BGC0002538 | 0.923188448 | A5252<br>(126) | A80510<br>(92) | A5858<br>(92) | A61715<br>(74) | T1312<br>(66) | A11345<br>(55) | A33995<br>(49) | A58051<br>(47) | A34001<br>(43) | A34053<br>(43) |
|  |  | BGC0001449 | 0.922354937 | A5252<br>(206) | A80510<br>(180) | A5858<br>(156) | A61715<br>(130) | A58051<br>(125) | A11345<br>(123) | A2056<br>(116) | A8567<br>(108) | A6562<br>(102) | A34001<br>(101) |

**Table 3.** Statistics of MIBiG version 3.1 and version 4.0.

| dataset | #BGCs | #compounds | #pairs | #BGCs<br>in pairs | #compounds<br>in pairs |
| --- | --- | --- | --- | --- | --- |
| version 3.1 | 2494 | 2688 | 3110 | 1906 | 2682 |
| version 4.0 | 2625 | 3000 | 3422 | 1987 | 2992 |
| only in 3.1 | 88 | 120 | 161 | 53 | 120 |
| only in 4.0 | 219 | 432 | 473 | 134 | 430 |

## Discussion

In this work, we present a generalizable framework for aligning biosynthetic gene clusters (BGCs) and their corresponding natural products within a shared latent space. By integrating large pretrained language models—BiGCARP for BGC sequences and MoLFormer for chemical structures— into a metric learning paradigm, our approach bridges genomic and chemical modalities through cross-modal representation learning. The resulting co-embedding space captures fine-grained relationships between protein domain architectures and molecular structures, enabling accurate retrieval of BGC-compound pairs and highlighting latent biochemical organization beyond sequence or structure alone.

Critically, the learned representations not only recapitulate known natural product classes but also support bidirectional retrieval tasks central to both traditional and inverse natural product discovery workflows. This dual capability allows for the prioritization of candidate biosynthetic clusters given a target molecule, or the identification of plausible chemical products from newly sequenced strains. As demonstrated across three distinct evaluation settings—including extrapolation to unseen compound classes and forward prediction of novel pairs from future database releases—the model exhibits strong performance and robustness. These results underscore its practical value in reducing the scale of experimental validation in high-throughput strain screening and retrosynthetic planning.

We compared our BGC-Chemical Co-Embedding (BCCoE) model against two non-aligned base-line methods: a standard nearest-neighbor (KNN) approach and a two-hop retrieval strategy (KNN-2hop). While KNN-2hop improves over KNN by incorporating indirect similarity chains, it consistently underperforms relative to BCCoE. Moreover, KNN-2hop exhibits greater sensitivity to the quality of the initial embeddings generated by foundation models. As shown in Figure 8, its performance degrades significantly when the similarity landscape becomes overly uniform—an issue that arises when foundation model embeddings are saturated (i.e., nearly all cosine similarities approach one). In contrast, BCCoE remains stable across diverse initialization conditions, suggesting that the alignment procedure provides meaningful inductive structure and regularization.

More broadly, the ability to align heterogeneous biological and chemical data types in a unified latent space introduces a flexible interface for downstream computational tasks. The structure of this space may be further exploited by generative models—such as diffusion or transformer-based decoders—to design novel small molecules conditioned on genetic inputs or, conversely, to propose synthetic gene clusters capable of producing specified target compounds. By grounding generative design in a metric-informed space trained on real-world biosynthetic constraints, this framework could serve as a foundation for automated retrosynthesis, combinatorial pathway engineering, or even closed-loop design of biosynthetic pathways.

## Methods and Materials

### Datasets

We used MIBiG version 3.1 and 4.0 (https://mibig.secondarymetabolites.org/download) in our study. Only active BGCs and their natural products are included. BGCs are represented by sequences of Pfam domains, which are generated from protein FASTA sequences of BGCs using HMMER version 3.4 and Pfam 31.0 model. Compounds are represented by their canonical SMILES. BGCs without Pfam domains or associated SMILES are excluded. The first two rows in Table 3 shows the number of active BGCs with Pfam sequences, the number of compounds with SMILES, BGC-compound pairs, number of BGCs in pairs and number of compounds in pairs in the two versions. The last two rows in Table 3 shows the number of BGCs, compounds and BGC-compound pairs in version 3.1 only or in version 4.0 only.

### BCCoE: BGC-chemical co-embedding for alignment

We use deep metric learning to map BGCs and compounds to a common co-embedding space to facilitate efficient bidirectional retrieval.

#### Problem statement

Formally, let ℬ be the space of BGCs and ℳ be the space of compounds. If a compound *m* ∈ ℳ is a natural product of a BGC *b* ∈ ℬ, then (*b, m*) is called a *positive pair*; otherwise (*b, m*) is called a *negative pair*. Let 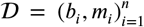 be a dataset of known positive BGC-compound pairs where *b* ∈ ℬ and *m*_*i*_ ∈ ℳ. The MIBiG database is one example of such a dataset. Pairs not in 𝒟 are either unknown positive pairs or negative pairs. Our goal is to discover new unknown positive pairs by using data in 𝒟 to learn the connections and linkage between BGCs and compounds. More specifically, we learn two functions *f* _ℬ_ : ℬ → ℝ^*d*^ and *f*_ℳ_ : ℳ → ℝ^*d*^ to map BGCs and compounds into a *d*-dimensional co-embedding space so that embeddings of a BGC and a compound from a positive pair are close to each other and embeddings of a BGC and a compound of a negative pair are farther apart. We hypothesize that BGCs and compounds part of the same (unknown) positive pairs should be close to each other in the co-embedding space as well, and given one element of an unknown positive pair, the other element can be efficiently retrieved using nearest neighbor search in the co-embedding space.

#### Model architecture

The overall architecture of our BCCoE model is shown in Figure 1 (A). BGC Pfam sequences and compound SMILES are converted to initial embeddings using a BGC foundation model and a chemical foundation model respectively. The initial embeddings of BGCs and compounds are in different embedding spaces, so they are not aligned as shown in Figure 1 (C). We then use a BGC Encoder and a chemical Encoder to map the initial embeddings to embeddings in a co-embedding space, and the mapped embeddings of BGCs and compounds from same positive pairs are aligned and close to each other in the co-embedding space as shown in Figure 1 (D).

We use BiGCARP (***Rios-Martinez et al., 2022***)(https://github.com/microsoft/bigcarp) to generate the initial 256-dimensional embeddings of BGCs and use MoLFormer (***Ross et al., 2022***)(https://huggingface.co/ibm/MoLFormer-XL-both-10pct) to get the initial 768-dimensional embeddings of compounds. Both foundation models are masked language models trained on unlabeled data using self-supervised learning. BiGCARP represents BGCs as chains of functional protein domains and uses a convolutional masked language model to learn meaningful representations of BGCs and their constituent Pfam domains. MoLFormer is a transformer encoder based chemical language model trained on over one billion SMILES strings. It leverages a linear attention mechanism and rotary positional embeddings for efficient training, and is able to accurately predict a diverse range of chemical properties. Other BGC or chemical foundation models can be employed in our model to generate the initial embeddings of BGCs and compounds as well, and we study two other foundation models in a later subsection.

The initial embeddings of BGCs and compounds are mapped to 64-dimensional embeddings in the co-embedding space via a BGC Encoder and a chemical Encoder respectively. The two encoders share the same model architecture as shown in Figure 1 (B) but they do not share model weights. Within the two encoders, the initial embeddings generated by foundation models are linearly transformed to 64-dimensional vectors first before they are passed through a two-layer transformer encoder. The output vector of the transformer encoder is averaged along the sequence dimension to remove the sequence dimension and is then concatenated with the mean vector of the initial embedding sequence. The concatenation of the two vectors is then passed through a batch normalization layer and a two-layer MLP to get the final 64-dimensional embeddings in the co-embedding space. The dimension of the feed-forward neural network in the transformer encoder and the MLP layer is set to 512. The length of the input sequences to the transformer encoder is set to 128 given that the sequence length of 92.6% of BGCs and 81.6% of compounds is no more than 128. If the sequence length of a BGC or a compound is larger than 128, we use the first length-128 sub-sequence only. We tested longer sequence length like 256 and it does not yield better performance than using 128.

After the embeddings of BGCs and compounds in the co-embedding space are generated, we can then use nearest neighbor search for efficient retrieval of BGCs that are likely to synthesize a query compound or retrieval of compounds that are likely natural products of a query BGC as illustrated in Figure 2. The similarity between a BGC *b* and a compound *m* in the co-embedding space is measured using cosine similarity defined in Equation 1, where *e*^*b*^ is the embedding of BGC *b, e*^*m*^ is the embedding of compound *m*, and *d*=64 is the dimension of the co-embedding space. The higher the similarity between a BGC and a compound is, the more likely the compound is a natural product of the BGC.

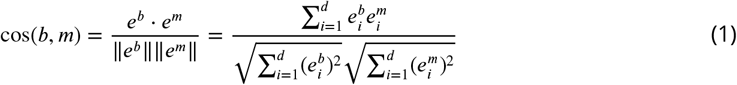

#### Model training

During training, we freeze the initial embeddings generated by foundation models to preserve their learned representations, and only update the parameters of the two encoders. This strategy allows us to leverage the rich feature representations learned by the two foundation models while reducing the risk of over-fitting.

To align the embeddings of BGCs and compounds in the co-embedding space, we employ metric learning with N-pair loss (***Sohn, 2016***) to maximize the similarity between positive BGC-compound pairs and minimize the similarity of negative pairs. In each batch, *N* BGC-compound positive pairs 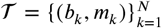 are randomly sampled from training data. Given a positive pair (*b, m*), N-pair loss with BGC *b* as the anchor is defined in Equation 2 and N-pair loss with compound *m* as the anchor is defined in Equation 3, where 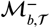 is the set of negative compounds of BGC *b* from 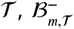 is the set of negative BGCs of compound *m* from 𝒯, and cos(*b, m*) is the cosine similarity defined in Equation 1. Note that negative BGCs of *m* and negative compounds of *b* are taken from other pairs in the same batch. A factor of 5 is multiplied to cos(*b, m*^−^) −cos(*b, m*) because it penalizes the difference more which leads to better performance than not multiplying in our experiments. The N-pair loss over batch 𝒯 is the mean of 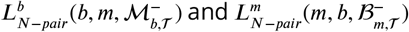 over all positive pairs in 𝒯 as given in Equation 4.

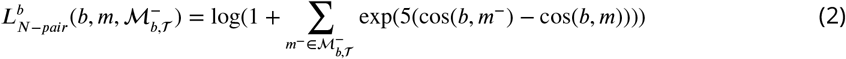

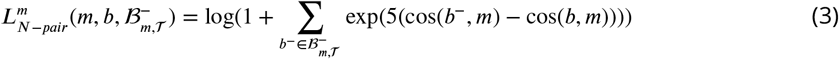

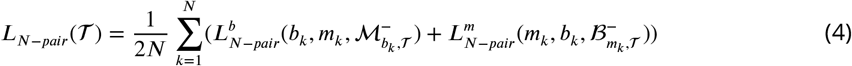

The model is trained using the Adam optimizer to minimize *L*_*N*−*pair*_(𝒯) in each batch. To achieve that, the optimizer needs to push negative pairs apart by reducing *cos*(*b, m*^−^) and *cos*(*b*^−^, *m*) and draw positive pairs close by increasing *cos*(*b, m*). Our model is trained using one cycle of cosine annealing schedule with an initial learning rate of 0.003 and a minimum learning rate of 0.0001. The number of epochs in one cycle is set to 100. The dropout rate is set to 0.1.

Besides using N-pair loss, we also tested two other loss functions for comparison: Binary Cross Entropy (BCE) loss and triplet loss. To calculate BCE loss over a bath 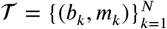, we randomly sample a negative batch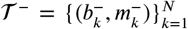 where 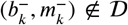. For each pair (*b, m*) in 𝒯 ∪ 𝒯 ^−^, we directly estimate the probability whether (*b, m*) is positive or not as given in Equation 5 below where *σ* is the sigmoid function. Here we multiply a factor of 5 to cos(*b, m*) to extend the value range of *ŷ* _*x*_ from [0.2689, 0.7311] to [0.0067, 0.9933].

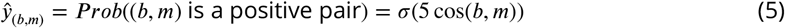

BCE loss over 𝒯 ∪ 𝒯 ^−^ is defined in Equation 6.

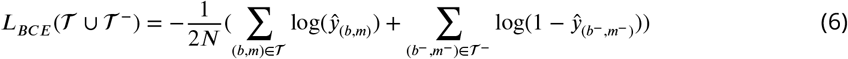

Triplet loss is similar to N-pair loss, but it is calculated on one negative pair only instead of N-1 negative pairs. Given a positive pair (*b, m*), triplet loss with BGC *b* as the anchor is defined in Equation 7 and triplet loss with compound *m* as the anchor is defined in Equation 8, where *b*^−^ and *m*^−^ are randomly sampled such that (*b*^−^, *m*) ∉ Q and (*b, m*^−^) ∉ Q. The triplet loss over batch 𝒯 is the mean of 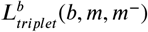 and 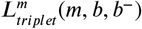 over all positive pairs in 𝒯 as given in Equation 9.

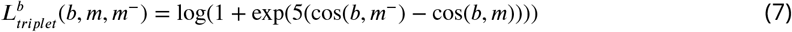

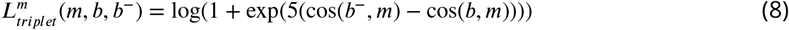

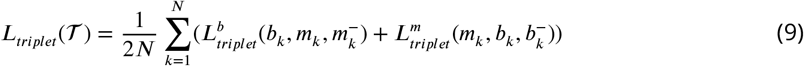

### Baseline: K-nearest neighbor search without alignment

We compare our BCCoE model with a more conventional approach of k-nearest neighbor search in the original embedding space directly without alignment. The assumption behind this baseline approach is that if two BGCs are very similar to each other, then they are likely to produce the same or similar natural products; and if two compounds are very similar to each other, then they are likely to be produced by the same or similar BGCs. We implemented two versions of this baseline, KNN and KNN-2hop, and both of them use cosine similarity over the original embeddings generated by BGC and chemical foundation models. The difference between the two is that KNN uses either similarities between BGCs or similarities between compounds depending on the query while KNN-2hop uses both similarities in order to retrieve novel BGCs or compounds not in the training data. Figure 7 (A) shows an example training data containing three positive BGC-compound pairs: (1, a), (2, b) and (3, b), where 1, 2, 3 are BGCs and a, b, c are compounds. Given a query BGC 4, the similarity between BGC 4 and the three BGCs in the training data are calculated over the initial BGC embeddings. Among the three BGCs in the training data, BGC 1 has the highest similarity of 0.9 with the query BGC, so its compound, compound a, is ranked first. KNN is unable to retrieve and rank compound c because compound c is not produced by any BGC in the training data. KNN-2hop overcomes this limitation by considering the similarities between compounds in addition to similarities between BGCs. In KNN-2hop, the route from the query BGC to a compound can be either through a neighbor BGC as in KNN or through a neighbor BGC (the first hop) plus a neighbor compound (the second hop). In the latter case, the final score is the product of the two similarities over the two hops. For both KNN and KNN-2hop, if there are multiple routes from a query BGC and a compound, the one with the highest score is used, and other routes are ignored. In the example in Figure 7, KNN-2hop is able to retrieve and rank compound c for the query BGC. Among the three routes from the query BGC to compound c, the one containing BGC 1 and compound a has the highest final score of 0.81, so this route is used and the other two routes are ignored. Figure 7 shows how compounds are retrieved and ranked for a given query BGC using KNN and KNN-2hop. The other direction of retrieving BGCs for a given query compound is done in a similar way.

**Figure 7.**
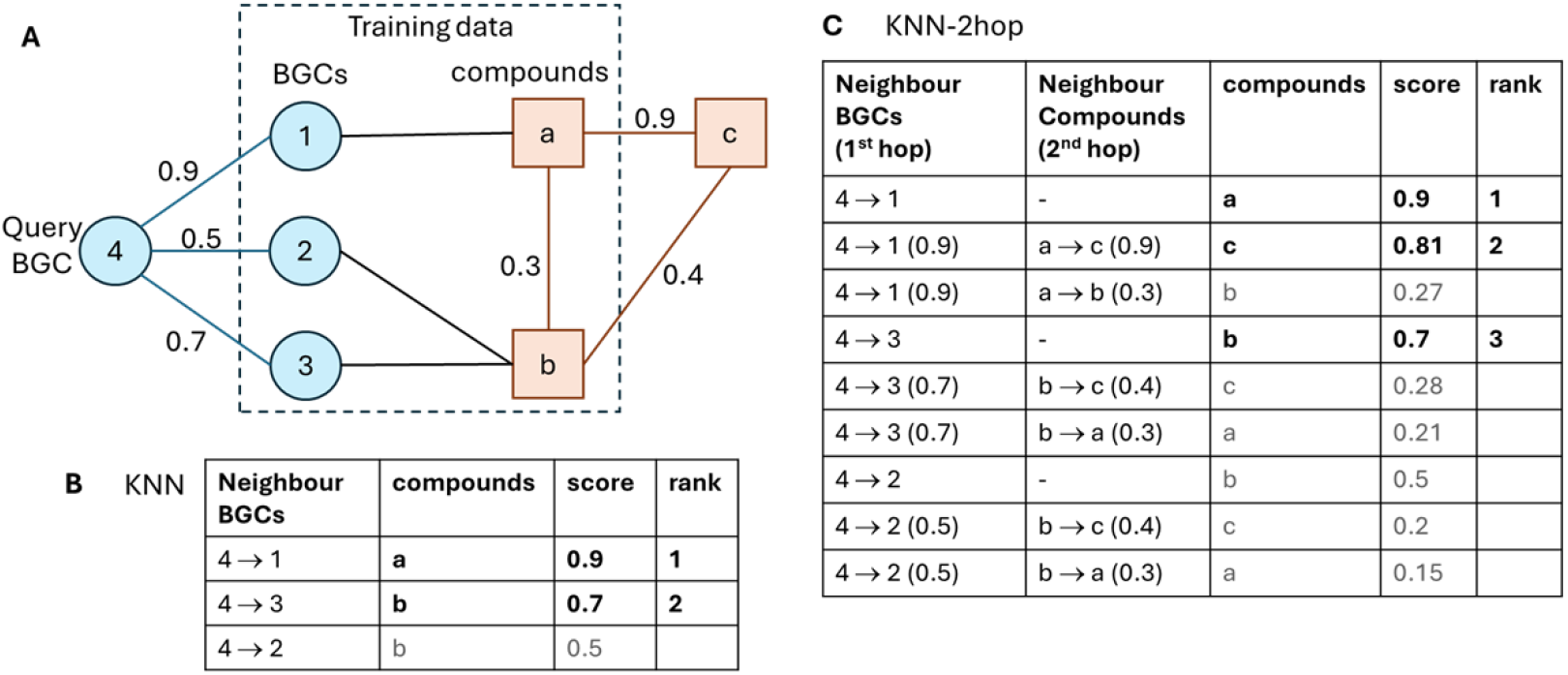
KNN and KNN-2hop. **(A)** An example training data. Similarities between the query BGC (BGC 4) and BGCs in the training data are calculated using the initial embeddings generated by a BGC foundation model. Similarities between compounds are calculated using the initial embeddings generated by a chemical foundation model. **(B)** compounds retrieved and ranked by KNN for the given query BGC. **(C)** compounds retrieved and ranked by KNN-2hop for the query BGC.

**Figure 8.**
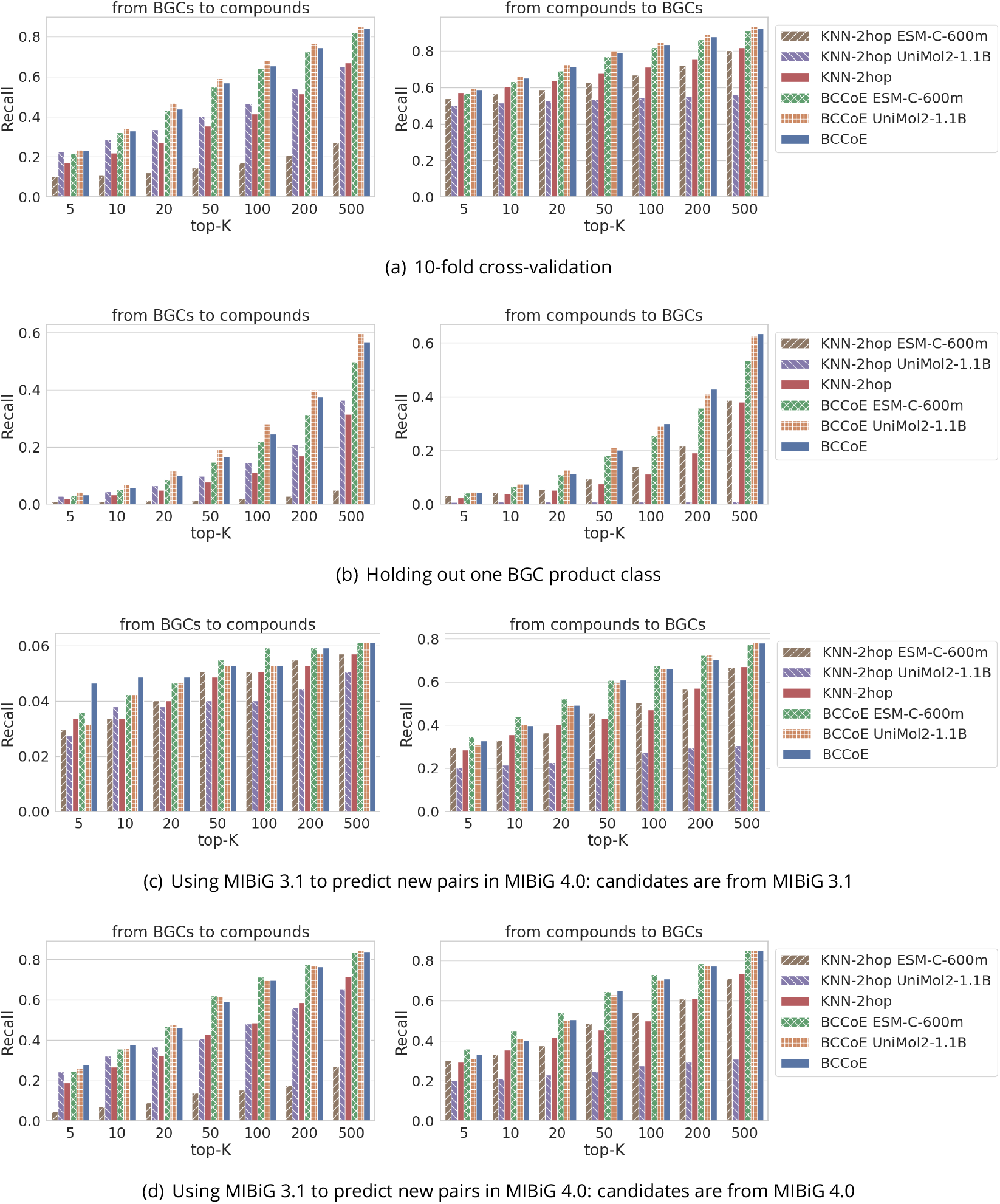
Recall of models for bidirectional retrieval when using different foundation models. Left: from BGCs to compounds; Right: from compounds to BGCs.

### Model evaluation

Our BCCoE model can be used for efficient bidirectional retrieval of BGCs and compounds to discovery novel positive BGC-compound pairs. For a given query BGC *b* and a list of candidate compounds, we can rank the compounds by their similarity to *b* in the co-embedding space. The most similar compounds are predicted as likely natural products of BGC *b*. Similarly, given a query compound *m* and a list of candidate BGCs, we rank the BGCs in descending order of their similarity to *m*. The top BGCs are predicted to produce compound *m*.

We evaluate our model using #hits, recall, precision and lift at top-K, which are commonly used to evaluate the performance of information retrieval algorithms. We split data in Q into training data and testing data. Training data is used to learn the embeddings of BGCs and compounds in the co-embedding space, and testing data is used as the ground-truth to calculate #hits, recall, precision and lift at top-K. If a BGC or a compound in the top-K actually occurs in the ground-truth set, then it is called a *hit*. Recall at top-K is defined as the number of hits among top-K predictions over the number of ground-truth pairs as given in Equation 10.

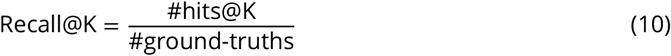

Precision at top-K is defined as the ratio of hits among top-K predictions as given in Equation 11, where N is the number of queries.

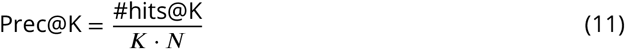

Lift is calculated by comparing the number of hits of a model with the number of hits of a baseline random guessing method as given in Equation 12. For the random guessing method, we use its average performance over 100 runs to ensure reliable measurements.

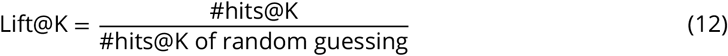

### Model performance for bidirectional retrieval

We studied the performance of our BCCoE model with N-pair loss and the two baseline KNN methods under several settings: 10-fold cross-validation, holding out one BGC product class and using MIBiG version 3.1 as training data to predict new pairs in MIBiG version 4.0.

#### 10-fold cross-validation

MIBiG 4.0 dataset is split randomly either based on BGCs or compounds into 10 folds. More specifically, if the task is to retrieve compounds for a given BGC, then data are split based on BGCs randomly so that a BGC and all its positive pairs are in one and only one fold (exp 1); if the task is to retrieve BGCs for a given compound, then data are split based on compounds randomly so that a compound and all its positive pairs are in one and only one fold (exp 2). Then each fold is used as testing data, and the remaining nine folds are used as training data to learn the model parameters.

This data-splitting strategy represents a relatively ideal situation for machine learning where the training data and the testing data have similar distributions as the data is split randomly. The performance of BCCoE, KNN and KNN-2hop under this setting are shown in Figure 3 and exp 1 & 2 in Table 1 where the performance of models is aggregated over the 10 folds. The BCCoE model shows much better performance than the two KNN methods when the task is to retrieve compounds for given BGCs. At top-10, KNN retrieves 12.9% of ground-truths with a precision of 2.23%; KNN-2hop retrieves 21.9% of ground-truths with a precision of 3.76%, which is a 68.8% improvement over KNN; BCCoE is able to pick up 32.9% of ground-truths with a precision of 5.66%, which is a 50.4% improvement over KNN-2hop. When the task is to retrieve BGCs for given compounds, both BCCoE and KNN-2hop perform very well. They can pick up 58.9% and 57.3% of the groundtruths respectively even at just top-5. In comparison, random guessing can only pick up 0.2% of the ground-truths at top-5.

#### Holding out one BGC product class

This experiment aims to test the ability of our model in retrieving compounds/BGCs with a novel product class. Table 4 shows the list of product classes in MIBiG 4.0, number of BGCs, number and percentage of positive pairs involving the BGCs, number of compounds, number and percentage of positive pairs involving the compounds in each product class. As BGCs or compounds can belong to more than one class, the sum of the percentages is more than 100%. We use each of these classes as a target class. If the task is to retrieve compounds for a given BGC, we split based on BGCs and use the BGCs of the target class and their positive pairs as the testing data (exp 3). If the task is to retrieve BGCs for a given compound, we split based on compounds and use compounds that are synthesized by BGCs of the target class and the positive pairs of the compounds as the testing data (exp 4). The remaining data are used as training data. For both splits, the training data do not contain any testing BGC/compound from the target class. The model only has access to the training data, so the target class is novel to the model.

**Table 4.** Statistics of BGC product classes.

| classes | #BGCs | BGC pairs |  | #compounds | compound pairs |  |
| --- | --- | --- | --- | --- | --- | --- |
|  |  | num | % |  | num | % |
| PKS | 932 | 1673 | 48.9% | 1483 | 1697 | 49.6% |
| NRPS | 737 | 1377 | 40.2% | 1242 | 1402 | 41.0% |
| ribosomal | 142 | 184 | 5.4% | 173 | 184 | 5.4% |
| saccharide | 141 | 202 | 5.9% | 152 | 227 | 6.6% |
| terpene | 152 | 292 | 8.5% | 263 | 307 | 9.0% |
| other | 335 | 592 | 17.3% | 520 | 611 | 17.9% |

**Table 5.** #new pairs with existing or new BGCs/compounds.

|  | existing BGCs | new BGCs | total |
| --- | --- | --- | --- |
| existing compounds | 25 | 4 | 29 |
| new compounds | 404 | 40 | 444 |
| total | 429 | 44 | 473 |

This data splitting strategy represents a very challenging situation for machine learning because the training data and the testing data have very distinct distributions. The performance of the three models under this setting is shown in Figure 4 and exp 3 & 4 in Table 1 where the performance of a model is aggregated over all the BGC product classes. Compared with using 10-fold crossvalidation, the performance of all the three models drops significantly in these two experiments as expected. Nevertheless, BCCoE achieves a recall of 5.9% and 7.5% at top-10 for the two retrieval tasks respectively, which is 74.5% and 89.2% better over the recall of 3.4% and 4.0% of KNN-2hop for the two tasks.

#### Using MIBiG 3.1 to predict new pairs in MIBiG 4.0

This setting represents a more realistic situation where we use known positive pairs to predict novel new pairs. There are 473 new BGC-compound pairs in MIBiG 4.0 compared with MIBiG 3.1 as shown in Table 3. We further divide these new pairs based on whether their BGCs and compounds exist in MIBiG 3.1 or are new in MIBiG 4.0 as shown in Table 5. The total number of unique BGCs in the new pairs is 196 and the total number of unique compounds in the new pairs is 457. If the task is to retrieve compounds for given BGCs, we use the 196 BGCs in the new pairs as query BGCs, and use either the compounds in MIBiG 3.1 (exp 5) or the compounds in MIBiG 4.0 (exp 7) as the candidate pool to retrieve compounds from. If the task is to retrieve BGCs for given compounds, we use the 457 compounds in the new pairs as query compounds, and use either the BGCs in MIBiG 3.1 (exp 6) or the BGCs in MIBiG 4.0 (exp 8) as the candidate pool to retrieve BGCs from. Some new pairs in MIBiG 4.0 contain new compounds or new BGCs not present in MIBiG 3.1, so in exp 5 and exp 6, algorithms will not be able to predict these new pairs because they use MIBiG 3.1 data only. In practice, augmenting the candidate set of BGCs or compounds (e.g. through prior knowledge) may help improve recall. In exp 7 and exp 8, we use the new BGCs and new compounds in ground-truths to expand the candidate pool, so they represent the upper bound of what we can achieve realistically when we expand the candidate pool. How to source new BGCs or compounds is beyond the scope of this paper, and we will study it in our future work.

The performance of the three models for predicting new pairs in MIBiG 4.0 is shown in Figure 5 and exp 5-8 in Table 1. Among the 473 new pairs, only 29 of them involving existing compounds in MIBiG 3.1. In exp 5 where compounds are retrieved from MIBiG 3.1, KNN-2hop can retrieve 16 out of the 29 new pairs at top-10 while BCCoE is able to pick up 23 at top-10, which is a 43.8% improvement. This number increases to 126 (26.6%) and 180 (38.1%) respectively for the two models when retrieving from compounds from MIBiG 4.0 (exp 7). Most of the new pairs involving existing BGCs. BCCoE is able to pick up 188 (39.7%) new pairs at top-10 when retrieving from existing BGCs from MIBiG 3.1 (exp 6).

In summary, BCCoE consistently outperforms KNN and KNN-2hop across all the tested scenarios with greater improvement when holding out one BGC product class, underscoring the model’s higher generalizability over the no-alignment approach.

### Performance of models using other foundation models

Besides BiGCARP and MolFormer, other foundation models can also be employed in our framework to generate the initial embeddings of BGCs and compounds. Here we test two other foundation models, ESM C (***Team, 2024***) and Uni-Mol2 (***Xiaohong et al., 2024***) to see how different foundation models affect the performance of BCCoE and KNN-2hop. ESM C is a language model trained on protein sequences. We use the 600M-parameter ESM C model to generate initial embedding sequences of BGCs. Uni-Mol2 is a molecular pretraining model integrating atomic level, graph level and geometry structure level features. We use the 1.1B-parameter Uni-Mol2 model to generate the initial embedding sequences of compounds. For both BCCoE and KNN-2hop, when we replace BiGCARP with ESM C, MolFormer is used to generate initial embeddings for compounds; when we replace MolFormer with Uni-Mol2, BiGCARP is used to generate the initial embeddings for BGCs.

The performance of BCCoE is relatively stable when different foundation models are used. For KNN-2hop, ESM C performs much worse than BiGCARP for retrieving compounds for BGCs (figures on the left) and Uni-Mol2 performs much worse than MolFormer for retrieving BGCs for compound (figures on the right). The reason behind this is that the rankings among compounds/BGCs retrieved by KNN-2hop using one hop or two hops depend on the distribution of the similarities. If the similarities over the first hop is all close to one, then compounds/BGCs retrieved using two hops will mostly be ranked at bottom. To better understand the behaviors of KNN-2hop, we show the distribution of cosine similarities over initial embeddings and embeddings in the co-embedding space in Figure 9. Similarities computed over initial embeddings generated by ESM C and Uni-Mol2 are both highly skewed with most of their values are close to 1, while similarities computed over initial embeddings generated by BiGCARP and MolFormer are more balanced. When ESM C is used and the task is to retrieve compounds for BGCs, similarities over the first hop are similarities between BGCs computed over initial embeddings generated by ESM C and 74.9% of them are above 0.85; similarities over the second hop are similarities between compounds computed over initial embeddings generated by MolFormer, and 96.1% of them are below 0.85. As a result, it is hard for compounds retrieved using two hops to make it to the top. The situation is even more extreme when Uni-Mol2 is used and the task is to retrieve BGCs for compounds. In this setting, similarities over the first hop are similarities between compounds computed over initial embeddings generated by Uni-Mol2 and 99.8% of them are above 0.9; similarities over the second hop are similarities between BGCs computed over initial embeddings generated by BiGCARP and 99.7% of them are below 0.9.

**Figure 9.**
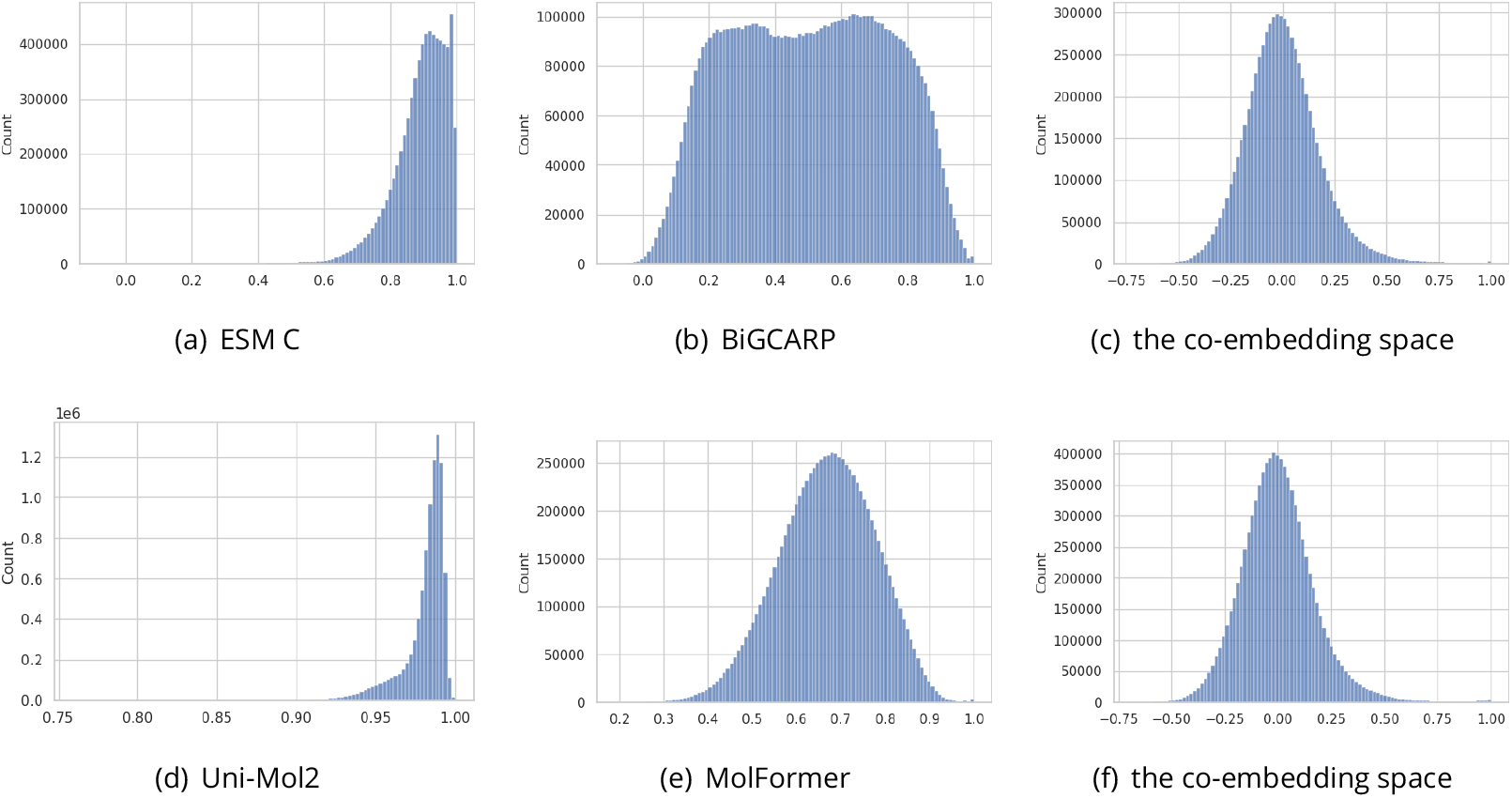
Distribution of similarities over initial embeddings and embeddings in the co-embedding space. Top row: similarities between BGCs; bottom row: similarities between compounds.

BCCoE does not have the above issue since it performs alignment in the co-embedding space first and then do retrieval in the co-embedding space. Similarities in the co-embedding space exhibit a bell-shaped normal distribution as shown in the last column of Figure 9. Being stable with respect to initial embeddings generated by different foundation models is another advantage of BCCoE over the no-alignment approach in addition to the higher performance in retrieval.

### Hyper-parameter tuning

In this subsection, we study the impact of several hyper-parameters, including loss function, sequence length, dimension of co-embedding space, factor multiplied to cosine similarity in Equation 2 and Equation 3, on the performance of BCCoE. We show Recall@K under 10-fold cross validation only in Figure 10. The results under other settings are similar.

**Figure 10.**
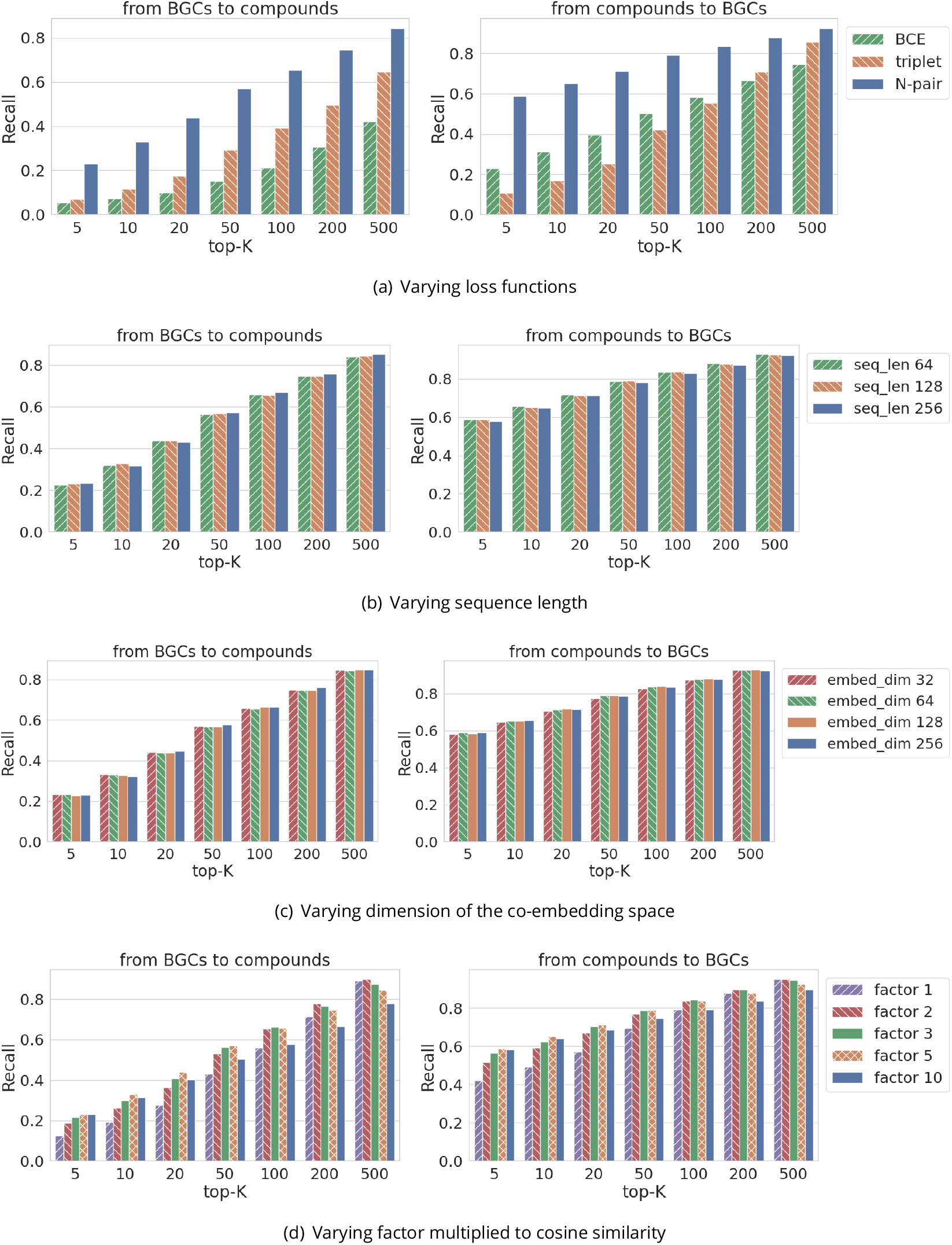
Recall of BCCoE for bidirectional retrieval when varying values of hyper-parameters.

Among the three loss functions, N-pair loss performs substantially better than BCE loss and triplet loss especially when K is small. BCCoE is not sensitive to sequence length and embedding dimension. For the factor multiplied to cosine similarity, larger values perform better when K is small and small values perform better when K is large. Considering that top predictions are more important, a value of 5 makes a good trade-off among different values.

The length of the input sequences to the transformer encoder in Figure 1 is set to 128. If the sequence length of a BGC or a compound is larger than 128, we use the first length-128 sub-sequence only. We tried another strategy which samples length-128 sub-sequences using a sliding window with a gap of 12 (10%) during inference, and then take the max over the samples as the final score. Figure 11 shows that the two strategies perform very closely.

**Figure 11.**
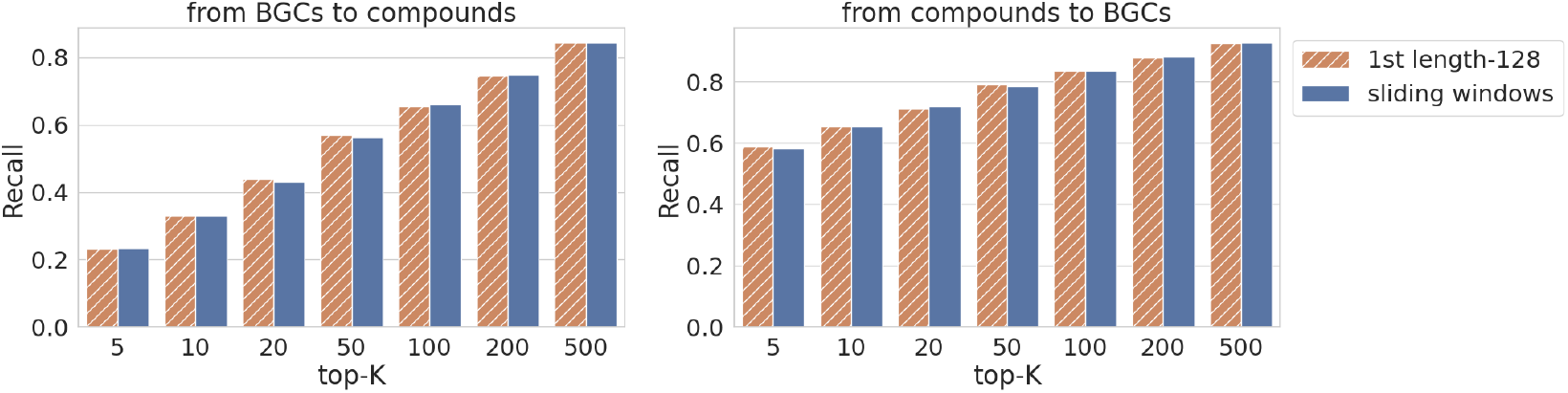
Recall of BCCoE for bidirectional retrieval with different sub-sequence sampling strategies.

### Top BGCs retrieved by BCCoE and KNN-2hop for new compounds in MIBiG 4.0

In this subsection, we take a closer look at the difference between the predictions made by BCCoE and KNN-2hop. We use both models to retrieve BGCs for new compounds in MIBiG 4.0 and BGCs from MIBiG 3.1 are used as the candidate pool. There are 19 ground-truth BGCs that are ranked by BCCoE within top-5 but are ranked by KNN-2hop beyond top-100, while there is only one groundtruth BGC that is ranked by KNN-2hop within top-5 but are ranked by BCCoE beyond top-100. These 20 cases are shown in Table 6. The routes from the query compounds to the BGCs retrieved by KNN-2hop are shown in Table 7. The differences demonstrate that even though KNN-2hop has inferior performance to BCCoE overall, but there are still cases where it can performs better than BCCoE. It may be beneficial to combine both approaches for even better performance. We will explore this in our future work.

**Table 6.** New BGC-compound pairs in MIBiG 4.0 that are ranked in top-5 by one model but are ranked beyond top-100 by the other model.

| no | query compounds | Hit BGCs | BCCoE |  | KNN-2hop |  |
| --- | --- | --- | --- | --- | --- | --- |
|  |  |  | score | rank | score | rank |
| 1 | hassallidin C | BGC0000369 | 0.581 | 1 | 0.851 | 142 |
| 2 | allocyclinone B | BGC0001500 | 0.716 | 1 | 0.868 | 137 |
| 3 | hassallidin D | BGC0000369 | 0.574 | 1 | 0.863 | 127 |
| 4 | 4-hydroxy-6-(2-oxoheptadecyl)pyran-2-one | BGC0002332 | 0.638 | 2 | 0.676 | 1541 |
| 5 | Isofuranonaphthoquinone A | BGC0001625 | 0.619 | 2 | 0.846 | 116 |
| 6 | Livipeptin | BGC0001168 | 0.606 | 3 | 0.669 | 1797 |
| 7 | paeninodin | BGC0001356 | 0.723 | 3 | 0.744 | 1297 |
| 8 | Adipostatin A | BGC0002332 | 0.728 | 3 | 0.706 | 1077 |
| 9 | dithioclapurine | BGC0001365 | 0.532 | 3 | 0.760 | 680 |
| 10 | tetrathioclapurine | BGC0001365 | 0.475 | 3 | 0.764 | 671 |
| 11 | trithioclapurine | BGC0001365 | 0.509 | 3 | 0.765 | 670 |
| 12 | reductasporine | BGC0001224 | 0.620 | 3 | 0.733 | 347 |
| 13 | 10-Oxodehydrodihydrobotrydial | BGC0000631 | 0.502 | 3 | 0.785 | 162 |
| 14 | 4-hydroxy-6-pentadecylpyran-2-one | BGC0002332 | 0.580 | 4 | 0.691 | 1219 |
| 15 | ent-kauren-16alpha-ol | BGC0002221 | 0.831 | 4 | 0.822 | 516 |
| 16 | microansamycin F | BGC0001666 | 0.411 | 4 | 0.860 | 126 |
| 17 | microansamycin E | BGC0001666 | 0.365 | 5 | 0.834 | 226 |
| 18 | fluostatin C | BGC0001904 | 0.710 | 5 | 0.842 | 118 |
| 19 | allocyclinone A | BGC0001500 | 0.652 | 5 | 0.837 | 109 |
| 1 | Secalonic acid B | BGC0001886 | 0.274 | 112 | 0.913 | 3 |

**Table 7.**
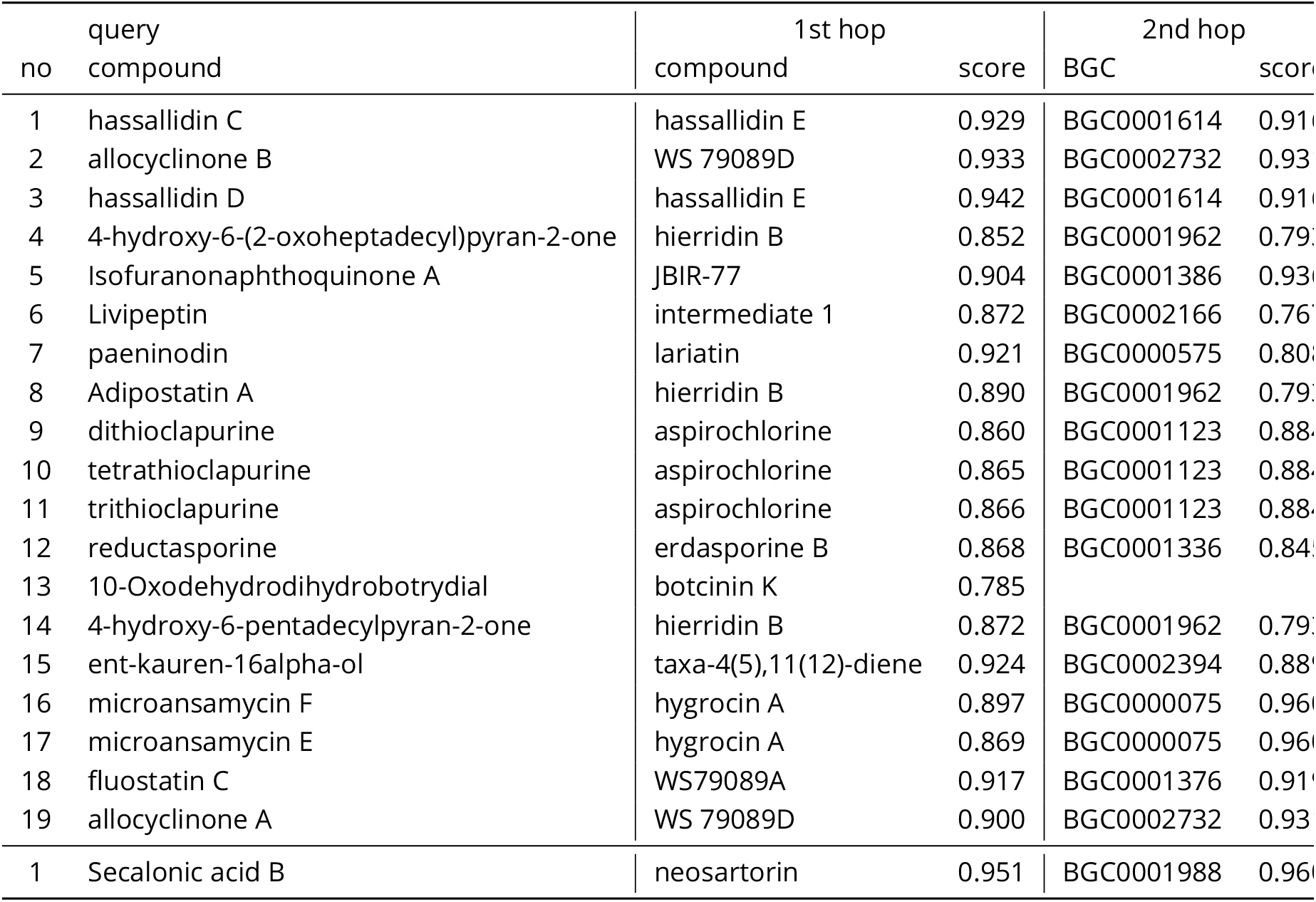
Routes from the query compound to the retrieved BGCs by KNN-2hop for the 20 cases in Table 6.

### Benchmarking retrieval of BGCs producing BE-54476-A/B

We applied Global Natural Products Social Molecular Networking (GNPS) (***Wang et al., 2016***), as described in the workflow in ***Tay et al. (2025***), to a publicly available dataset comprising 54 microbial strains from an independent study (***Tay et al., 2024a***), each with paired liquid chromatography-mass spectrometry (LC-MS) data and whole genome sequences. Our MS-based search identified one strain, A58051, producing detectable quantities of BE-54476-A and BE-54476-B.

BCCoE and KNN-2hop were used to identify the top 5 MIBiG 4.0 BGCs producing BE-54476A and BE-54476B. For each BGC ***B***_*i*_, composed of *n* genes {*g*_1_, *g*_2_, …, *g*_*n*_}, we extracted the corresponding amino acid sequences and searched each gene independently against the whole-genome sequences of the 54 microbial strains using tblastn ((***Altschul et al., 1997***), (***Camacho et al., 2009***)), with the BLOSUM62 substitution matrix and an E-value cutoff of *E <* 10. Let *m*_*ij*_ denote the number of significant tblastn alignments for gene *g* _*j*_ in strain ***S***. The total hit score 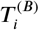 for strain *S* _*i*_ with respect to BGC ***B*** is defined as:

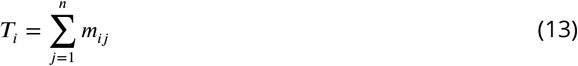

Strains were ranked in descending order of 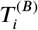, and the top 10 strains for each BGC were retained.

## Acknowledgments

This work was supported by RIE2025 A*STAR Central Funds - Strategic Programme Funds (SIBER 2.0, Project No. C2333017001). GO is a recipient of funding from the A*STAR Graduate Academy under the National Science Scholarship (BS-PhD).

Appendix 1

## Notes

### Competing Interest Statement

The authors have declared no competing interest.

